# Ser-653-Asn substitution in the acetohydroxyacid synthase gene confers resistance in weedy rice to imidazolinone herbicides imazapic and imazapyr in Malaysia

**DOI:** 10.1101/2019.12.19.882159

**Authors:** Rabiatuladawiyah Ruzmi, M. S. Ahmad-Hamdani, Norida Mazlan

## Abstract

The IMI-herbicides rice package has been recognized by all means among the most efficient chemical approaches for weedy rice control nowadays. Inevitably, the continuous and sole dependence, as well as ignorance on the appropriate use of imidazolinone herbicides in the IMI-herbicides rice package by rice growers has caused the development of herbicide resistance in weedy rice populations across many IMI-herbicides rice package adopted countries, inclusive of Malaysia. Hence, a comprehensive study was conducted to elucidate the occurrence, level, and mechanisms endowing resistance to IMI-herbicides on field-reported resistant (R) weedy rice populations collected from IMI-rice fields in Kampung Simpang Sanglang, Perlis (A), Kampung Behor Mentalon, Perlis (B), and Kampung Sungai Kering, Kedah (C). The collected weedy rice populations were compared with a susceptible weedy rice population (S), an imidazolinone-resistant rice cultivar (IMI-rice), and a susceptible local rice cultivar (MR219). Dose-response experiments were carried out using commercial IMI-herbicides (premix of imazapic and imazapyr) available in the IMI-herbicides rice package, in the seed bioassay and whole-plant dose-response. Based on the Resistance Index (RI) quantification in both experiments, the cross-resistance pattern of weedy rice populations and rice varieties to imazapic and imazapyr was determined. Molecular investigation was carried out by comparing acetohydroxyacid synthase (AHAS) gene sequences between resistant (R) weedy rice populations (A, B, and C), S population, IMI-rice, and MR219. Evidently, the AHAS gene sequences of R weedy rice were identical to the IMI-rice, revealing that amino acid substitution of Ser-653-Asn occurs in both R populations and IMI-rice, but neither in MR219 nor S plants. *In vitro* assays were conducted using analytical grade imidazolinone herbicides of imazapic (99.3%) and imazapyr (99.6%) with seven concentrations. The results demonstrated that the AHAS enzyme extracted from R populations and IMI-rice were less sensitive to IMI-herbicides in comparison to S and MR219, further supporting the IMI-herbicides resistance was conferred by target site mutation. In conclusion, the basis of imidazolinone resistance in selected populations of Malaysia weedy rice was due to a Ser-653-Asn mutation that reduced sensitivity of the target site to IMI-herbicides. The current study presents the first report of resistance mechanism in weedy rice in Malaysian rice fields.

## Introduction

Resistance to herbicides is commonly referred to the ability of the plants to inheritably survive and reproduce after the exposure to herbicide application with a spray rate that normally kills the wild/susceptible type [1,2]. Resistance in plants may naturally occur or may be induced by either genetic engineering or mutagenesis [3]. Two types of herbicide resistance mechanisms in plants have been debated; target site and non-target site mechanisms [4]. In target site mechanism, the target enzymes experience reduction in sensitivity to the herbicide due to mutation(s) at the site of action that prevent the herbicide from binding to the enzyme, and in the latter overexpression of the enzyme such as gene amplification or changes in gene promoter was also reported. In contrast, non-target site resistance mechanism happened with the presence of any one of this factors or the combination of the mechanisms which are the rapid herbicide detoxification, herbicide sequestration, decreased in herbicide penetration rate, and reduced herbicide translocation [4,5].

Acetohydroxyacid synthase (AHAS, EC 2.2.1.6) is a biosynthetic enzyme group that catalyzes the *de novo* step in synthesizing branched-chain amino acids, *viz* valine, leucine, and isoleucine. These amino acids are building blocks of protein, essential for the development and growth of new cells in plants, and their absence inhibits the growth of plant shoot and root, ultimately leading to plant death [6]. Acetohydroxyacid synthase (AHAS) enzyme is the target of commercial herbicides including imidazolinone and more than 50 AHAS-targeting herbicides are currently used commercially in the world [7].

The herbicide-tolerant crop, particularly IMI-herbicides rice package (e.g. Clearfield^®^ Rice Production System or CPS) has become among the preferred effective chemical-based methods to control weedy rice problems in Malaysia because previous chemical implementation in controlling weedy rice was not possible due to its genetic similarity with the conventional cultivated rice varieties. The imidazolinone AHAS inhibitors imazapic and imazapyr were chosen in the development of the IMI-herbicide-resistant rice technology to control weedy rice in Malaysia. In Malaysia, the imidazolinone (IMI)-resistant rice (IMI-rice) is the only herbicide-resistant rice that has been commercialized hitherto. The introduction of the non-transgenic imidazolinone-tolerant rice varieties (IMI-rice) package in Malaysian rice fields in late 2010 had received overwhelming response, and to the extent of continuous adoption of the IMI-herbicides rice package by rice growers. Subsequently, several years after the introduction, many weedy rice populations have been reported failed to be effectively controlled by the IMI-herbicides. This phenomenon has greatly affected the staple food industry in Malaysia, further leaving a question whether herbicide resistance is starting to develop in these weedy rice populations.

Weedy rice (*Oryza sativa* L. complex) has continuously become a long unsettled weed problem in the Malaysian rice granaries since the establishment of direct seeding method in the early 1980s [8]. It is closely related to cultivated rice, making the use of selective herbicides to control weedy rice is limited [9]. The use of herbicide-resistant rice cultivars such as IMI-herbicides rice package, advocated by researchers and government in 2010 has been a successful strategy for selective control of weedy rice in cultivated rice. Nonetheless, similar to other IMI-herbicide rice packages in other countries, over-reliance on this weedy rice management approach by rice growers has taken its toll, with an escalating field-reports on the reduced effectiveness of IMI-herbicides rice package to control weedy rice [10]. Another concern is on the possibility of hybridization to occur between IMI-rice and weedy rice, and the potential crosses among the weedy rice variants, which might increase its weediness, and those impact to the future of herbicide-resistant technology in rice [11], although this theory has yet to be proved in Malaysia rice granaries. Concomitantly, mutations in the AHAS gene, which conferred resistance to imazethapyr, have been reported in weedy rice that was growing in imazethapyr CPS rice fields in Arkansas, USA [3,12].

Currently, weedy rice has become the eighth resistant weed species in Malaysia’s rice fields [13] after been recognized as the most problematic weed in almost all rice granaries in Malaysia [14]. Recently, rice-growers who have been practicing IMI-herbicides rice package for more than eight planting seasons in Selangor, Perak, Pulau Pinang, Kedah, and Perlis of Malaysia have been complaining about the reduced efficacy of IMI-herbicides to control weedy rice and other weed species in their fields. In more recent findings, it was reported that weedy rice population collected from Pendang, Kedah have been confirmed to have developed resistance to the IMI-herbicides with the resistance level up to 67-fold greater than the susceptible population [15]. The risk of resistance evolution in weedy rice to the IMI-herbicides in Malaysia is now highly alarmed. Thus, the objectives of the present work were (1) to quantify the occurrence and level of resistance to AHAS IMI-herbicides in putative resistant weedy rice populations, (2) to identify the AHAS gene mutations conferring target site resistance mechanism to AHAS IMI-herbicides in the confirmed-resistant weedy rice populations, and (3) to investigate the existence of possible non-target site resistance mechanisms in the resistant weedy rice individuals by comparing AHAS enzyme activity and inhibition by IMI-herbicides.

## Materials and methods

### Seed source

Suspected resistant (R) weedy rice populations were collected from IMI-herbicides rice fields in three townships in the border of Kedah and Perlis states of Malaysia; Kampung Simpang Sanglang, Perlis (hereafter referred to as population A), Kampung Behor Mentalon, Perlis (population B) and Kampung Sungai Kering, Kedah (population C). Meanwhile the known herbicide susceptible (S) population seeds were obtained from Kampung Tanjong, Kedah from the fields that never been exposed to IMI-herbicides previously. The IMI-rice variety (MR220CL2) and a non IMI-rice variety (MR219) were used as positive control and the seeds were obtained from Department of Agriculture Malaysia (DOA). The sampling locations for suspected R were chosen based on the complaint received from the rice growers regarding the ineffectiveness of IMI-herbicides rice package to control weedy rice in their rice fields, plus Kedah and Perlis states are among the pioneers to adopt IMI-herbicides rice package following its launch in 2010. The seeds survey was conducted following the zig-zag pattern during the first planting season of the year e.g. February, 2017. Seeds were collected at around one week before rice harvesting operation took place. The collected seeds were placed in paper bag and brought immediately to the laboratory. Seeds were manually threshed, cleaned, and sorted to remove contaminants. Then, seeds were air-dried until the moisture content reached below 14%. Lastly, the seeds were then labeled accordingly, sealed in a plastic bag and stored in a refrigerator at 4°C until use.

### Seed bioassay

Seeds from each R and S weedy rice populations were treated with seed growth enhancer (Zappa™) for 24 hours (h) to induce seed germination, drained and incubated at room temperature (24-26 °C) until the radicle emerged for about 1 mm. Twenty uniform pre-germinated seeds were placed in 9-cm-diameter plastic disposable petri dishes, containing two sheets of Whatman No. 1 filter paper, and aliquots of 5 mL of aqueous commercial IMI-herbicides (OnDuty™ WG, BASF Malaysia Sdn. Bhd.) solutions together with the Tenagam surfactant at seven different concentrations; 0 (distilled water only), 0.2, 0.4, 0.77, 1.6, 3.1 and 6.2 mg/L (where 0.77 mg/L represents the recommended rate, RR). The dishes were transferred to a growth chamber and incubated at 30/20 °C (day/night temperature) with a 12 h/12 h (day/night cycle), under fluorescent light (8500 lux) with completely randomized design. Relative humidity ranged between 30% and 50%. The lids of the petri dishes were not sealed to allow gas exchange and to avoid an anaerobic condition. Five milliliters of distilled water were added daily, beginning from 24 h after IMI-herbicides application into the petri dishes to maintain the moisture. Fourteen days after treatment (DAT), the germinated seedlings were counted and shoot and root length were measured from the point of attachment of the seed to the tip of the epicotyl and hypocotyl, respectively. Experiment was repeated twice.

### Whole-plant dose response

The whole-plant dose response experiment was conducted at glasshouses compound in Ladang 2, Universiti Putra Malaysia during the period from March to August 2017. Seeds from R and S weedy rice populations were treated with seed growth enhancer, Zappa™ for 24 h, drained, and incubated at room temperature until the seeds began to germinate. Ten emerged seedlings of weedy rice (about 1-2 cm) were transferred to 37 cm × 30 cm × 7.5 cm plastic trays containing commercial paddy soil at 2/3 full and the plants were grown in trays with continuous 2-5cm flooded condition during the experimental period. Trays were arranged in a glasshouse with a day/night temperature of 33/22°C. The regular maintenance of the plants was handled following ‘Manual Penanaman Padi Lestari’ by the Malaysian Agricultural Research and Development Institute (MARDI). At the 1-2-leaf stage, all weedy rice and rice varieties seedlings were treated with commercial IMI-herbicides imazapic+imazapyr (OnDuty™ WG, BASF Malaysia Sdn. Bhd.) mix with Tenagam surfactant at seven different rates; 0 (water only), 38.5, 77.0, 154.0, 308.0, 616.0, 1232.0 g a.i./ha (where 77.0 g a.i/ha represents the recommended field dose, RR). Herbicides treatment were applied using a compression type sprayer with detachable flat fan nozzle, delivering 200 L/ha at a spray pressure of 150 kPa. The average temperature during the experiment was almost similar to the actual condition in rice fields, ranging from 22 to 33°C. At 21 days after herbicides treatment, the survived plants were counted and visual assessments on plant survival were made. Plants that survived and continuously produced new shoots/tillers following herbicide treatment were regarded as confirmed R and plants displayed severe symptoms of leaf chlorosis, desiccation, retarded growth or no new active growth and eventual plant death were considered as S. The rice varieties that act as a positive control was assessed in similar manner. Both treated and untreated plants were cut at 1 cm aboveground, dried at 65 °C for 72 h, and weighed. The dry weight of all plants (dead and alive) was recorded for each population. Plants surviving IMI-herbicides application from recommended rate and above, as well as the shoot materials from untreated S, IMI-rice, and MR219 plants were allowed to regrow, produce new shoots and leaf sample were taken individually and stored at −80°C for AHAS gene mutation study. Experiment was repeated twice.

### DNA extraction and amplification of AHAS gene fragment

The post-harvest shoot materials from the R weedy rice populations (populations A, B, and C) and the untreated plants of weedy rice S, IMI-rice, and MR219 were sampled individually for genomic DNA extraction and gene sequencing experiment for AHAS gene mutation study. Fifty-two leaf samples from R weedy rice plants and three from S weedy rice, IMI-rice, and MR219 were each taken approximately 100 mg per plant and extracted using a modified cetyltrimethylammonium bromide (CTAB) protocol [16]. The nucleotide sequence of AHAS gene fragment covering potential mutation sites was amplified by polymerase chain reaction (PCR) using forward primer 5’-GTAAGAACCACCAGCGACACC-3’ and reverse primer 5’-GATGCATATGCCTACGGAAAAC-3’ [17] to study the molecular basis of resistance in weedy rice populations and rice varieties to IMI-herbicides. The 15 μl PCR cocktail consisted of 7.5 μl of PCR MyTaq Red Mix (Bioline, GmbH, Germany), 1.0 μl of genomic DNA template (70 ng/μl), 1 μl of each primer (10 mM), and 4.5 μl of deionized water. The PCR was carried out in a heated lid PCR machine Thermal Cycler T100 (Bio-Rad, Hercules, CA, USA) using the following cycle: 95°C for 5 min for denature, then 34 cycles of 30 s at 95°C, anneal at 67°C for 30 s, 120 s of elongation at 72°C, followed by final elongation step at 72°C for 5 min. The amplified PCR products were resolved on 1.2% agarose gels stained with FloroSafe DNA Stain (Axil Scientific Pte Ltd, Singapore) by using 1kb DNA ladder (Fermentas, Thermo Scientific) as a reference. Unpurified PCR products were sent immediately to commercial sequencing service (First BASE Laboratories Sdn. Bhd.) for purification, gel extraction by general agarose, and DNA sequencing. The AHAS sequence chromatograms for all samples were visualized using DNA Baser Assembler v5.15.0. The same software was used to align and compare the obtained nucleotide sequences, which then translated to respective amino acid sequence using BLASTX (http://www.ncbi.nlm.nih.gov/). All amino acid sequences were standardized to the *Arabidopsis* sequence (NM114714).

### AHAS enzyme inhibition assay

All IMI-herbicides treated parent plants of R weedy rice populations, and untreated S and rice varieties taken for gene mutation study as mentioned above were allowed to regrow for seed production. Seeds were harvested and the progeny lines of R, S, and rice varieties were regrown again in the glasshouse until they reached 3-4-leaf stage. Leaf tissue from progeny lines of R and S populations, and the rice varieties were used for *in vitro* AHAS activity assay. The AHAS *in vitro* activity and inhibition assay was conducted according to the method described by Yu et al. [18] with some modifications. The protein concentration of the extract was determined by spectrophotometer using Bradford method and the extraction was instantly used in the inhibition assay. Certified analytical standards for IMI-herbicides of imazapic (99.3%) and imazapyr (99.6%) were incubated with partially purified enzyme extracts for 60 min, and the reaction was stopped by addition of 40 μl of 6NH_2_SO_4_ then incubated at 60°C for 15 min to convert acetolactate to acetoin. Acetoin formed from acetolactate was quantified colorimetrically (530 nm). Each enzyme assay was performed with three independent extractions and protein concentration was measured four times in each sample.

### Statistical analysis

The petri dishes were arranged in completely randomized design for seed bioassay and the experimental trays were laid out in a randomized complete block design in the whole-plant dose response experiment with four replications per treatment for both experiments. Seed germination rate, shoot length, root length, plant survival rate and shoot dry weight were expressed as a percentage of the untreated control for that population. All analyses were conducted using Sigmaplot Version 11.0 (Systat Software Inc., GmbH, Germany). A non-linear regression using the logistic response equation (Equation 1) proposed by Knezevic et al. [19] was used to obtain the dose response curve. Similarly, data obtained from AHAS enzyme inhibition assay were expressed as the percentage of untreated controls and subjected the same analysis as in seed bioassay and whole-plant dose response experiment. Equation 1 is as below:

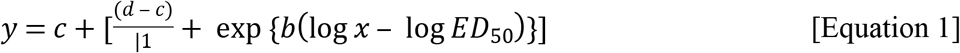

where,

c is the lower limit

d is the upper limit

b is the slope and

ED_50_ is the dose required to give 50% effect.

In the regression equation, the dose of herbicide was the independent variable (x), and the percentage of the control (plant survival) was the dependent variable (y). The fitted equations were used to calculate the amount of herbicide exhibiting a 50% reduction in dry weight of plant (GR_50_ value), 50% plant survival reduction (LD_50_ value) or 50% inhibition of *in vitro* AHAS activity (I_50_).

## Results

### Seed bioassay

The IMI-herbicides resistance level in the weedy rice populations in this experiment were quantified based on the Resistance Index (RI) value of herbicide rate resulting in 50% mortality (LD_50_). The resistance level to IMI-herbicides has been categorized as high (> 15), moderate (7 to 15), low (≥2 to 6) and sensitive (< 2), comparatively to Iwakami et al. [20] and Merotto et al. [21]. Table 1 shows the RI values of each population of weedy rice and rice varieties based on the quantification of seed germination percentage in seed bioassay. The IMI-rice variety yielded the highest RI value, followed by populations A and B at moderate level. Meanwhile S population and MR219 shared a similar sensitivity to the IMI-herbicides at seed germinating stage.

**Table 1.** Analysis of log-logistic of the IMI-herbicides dose needed to exhibit 50% mortality (LD_50_) and the resistance index (RI) of weedy rice populations and rice varieties in seed bioassay. Standard errors are in parentheses.

| Populations | LD<sub>50</sub> (mg/L) | RI<br>(LD<sub>50</sub> R/LD<sub>50</sub> S) | Resistance level |
| --- | --- | --- | --- |
| A | 2.25 (0.56) | 16.35 | High |
| B | 2.05 (7.31) | 14.92 | Moderate |
| C | 0.18 (0) | 1.30 | Sensitive |
| S | 0.14 (0) | - | Sensitive |
| IMI-rice | 7.07 (0.15) | 50.5 | High |
| MR219 | 0.14 (0) | 1.07 | Sensitive |

With increasing IMI-herbicides concentration, all populations experienced greater reduction in germination rate, and shorter shoots and roots length as compared to their untreated controls. The recommended rate of IMI-herbicides used in this experiment was equal to 0.77 mg/L. Population A, population B, and IMI-rice that survived until 4-fold the recommended concentration, nonetheless experienced complete germination inhibition at the concentration of 6.16 mg/L which was 8 times higher than the recommended concentration (Fig 1). Meanwhile for population C, population S and MR219, the pre-germinated seeds were totally controlled at only half recommended concentration (0.39 mg/L).

**Fig 1.**
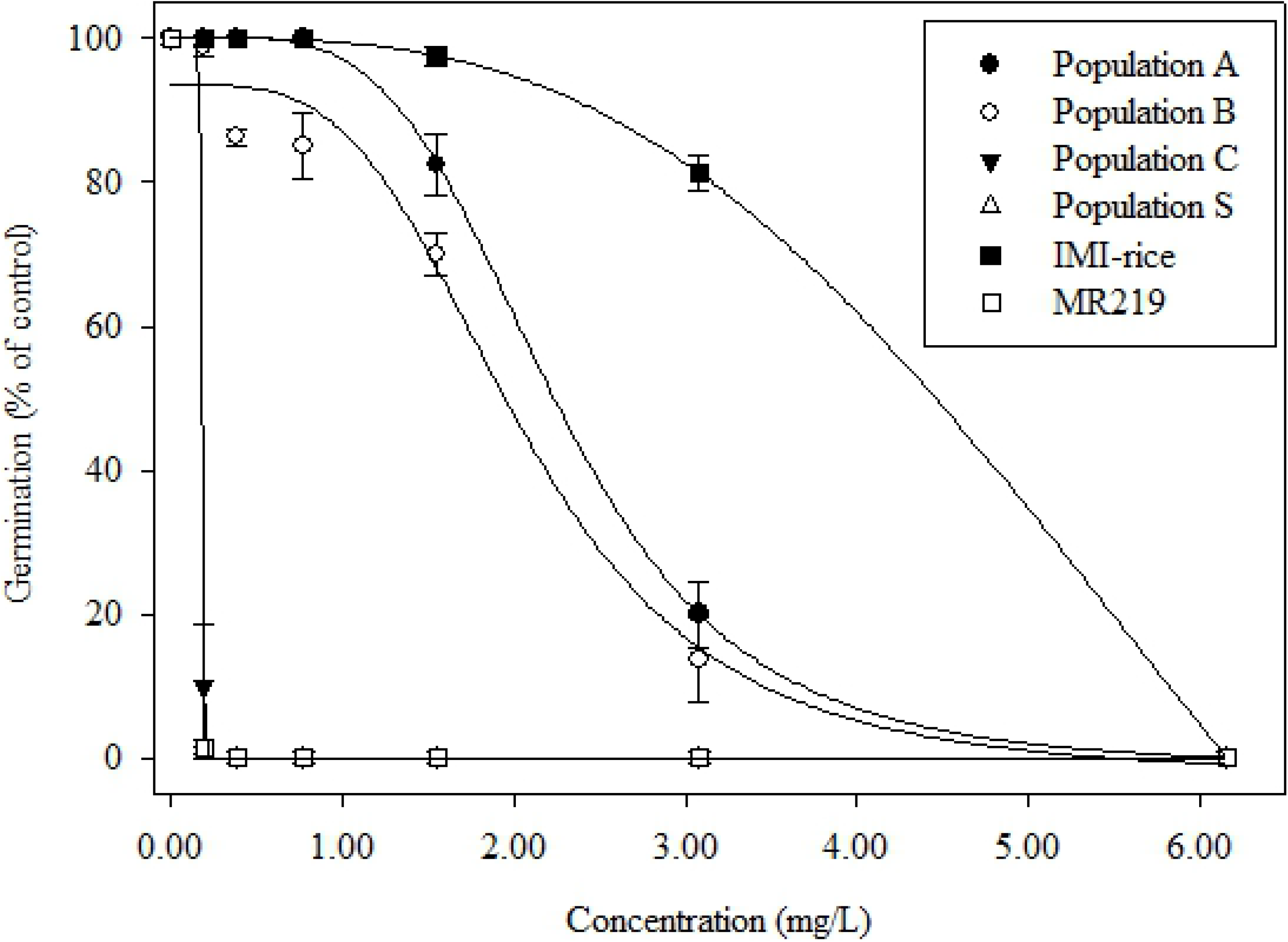
Germination rate (% of control) of weedy rice populations and rice varieties 14 days after treatment in seed bioassay.

Figs 1-3 displays the germination rate (% of control), shoot length (% of control) and root length (% of control) of all weedy rice populations and rice varieties, respectively 14 days after treatment (DAT) in seed bioassay. The IMI-herbicides effect on shoot and root length of R and S weedy rice populations and rice varieties was visible as soon as germination initiated, and lasted until 14 days of incubation. The treated seeds showed severe injury symptoms e.g. stunted epicotyl and hypocotyl growth, which led to inhibition of shoot and root elongation when compared to the untreated control seeds (Figs 4 and 5)

**Fig 2.**
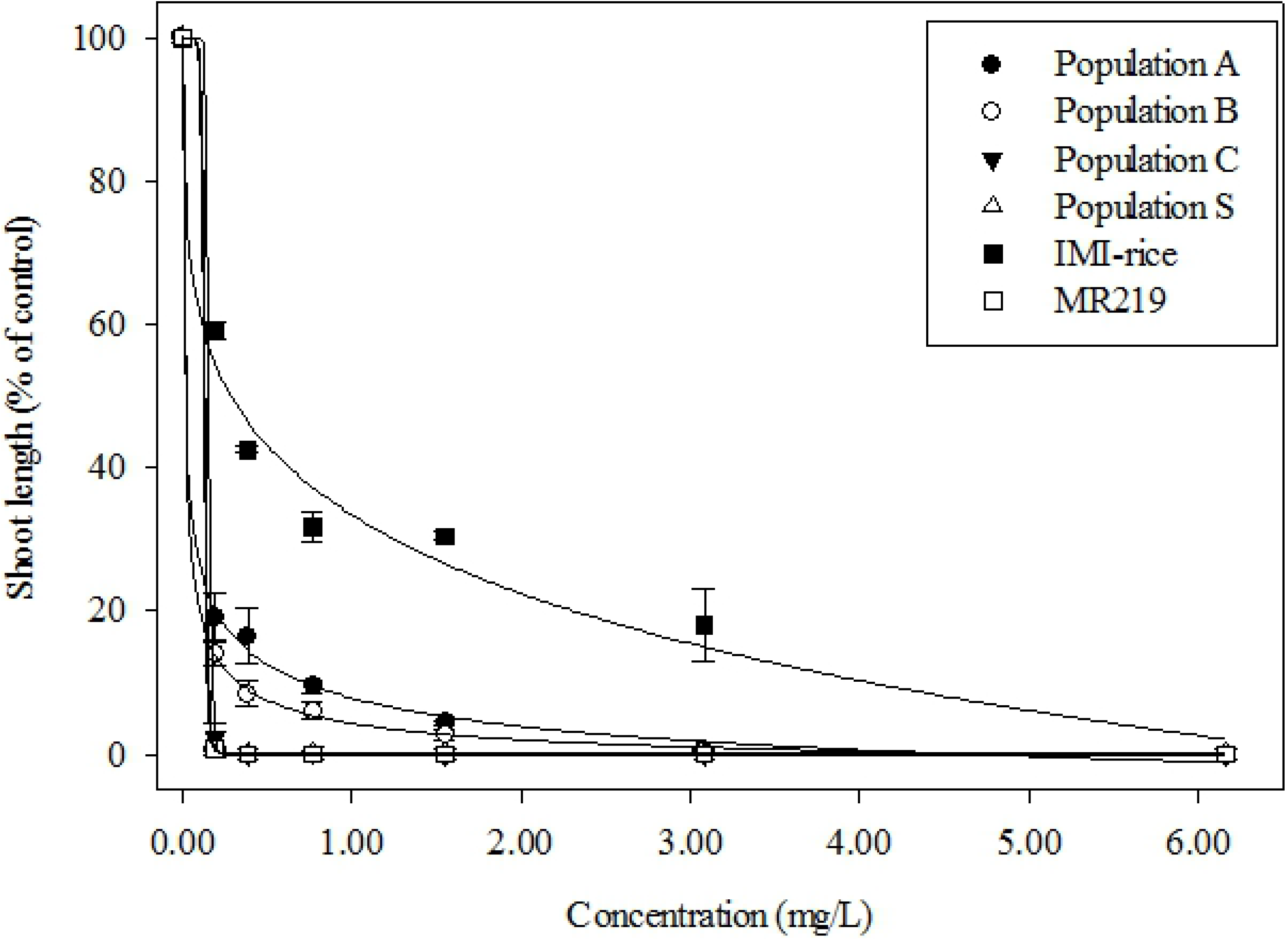
Shoot length (% of control) of weedy rice populations and rice varieties 14 days after treatment in seed bioassay.

**Fig 3.**
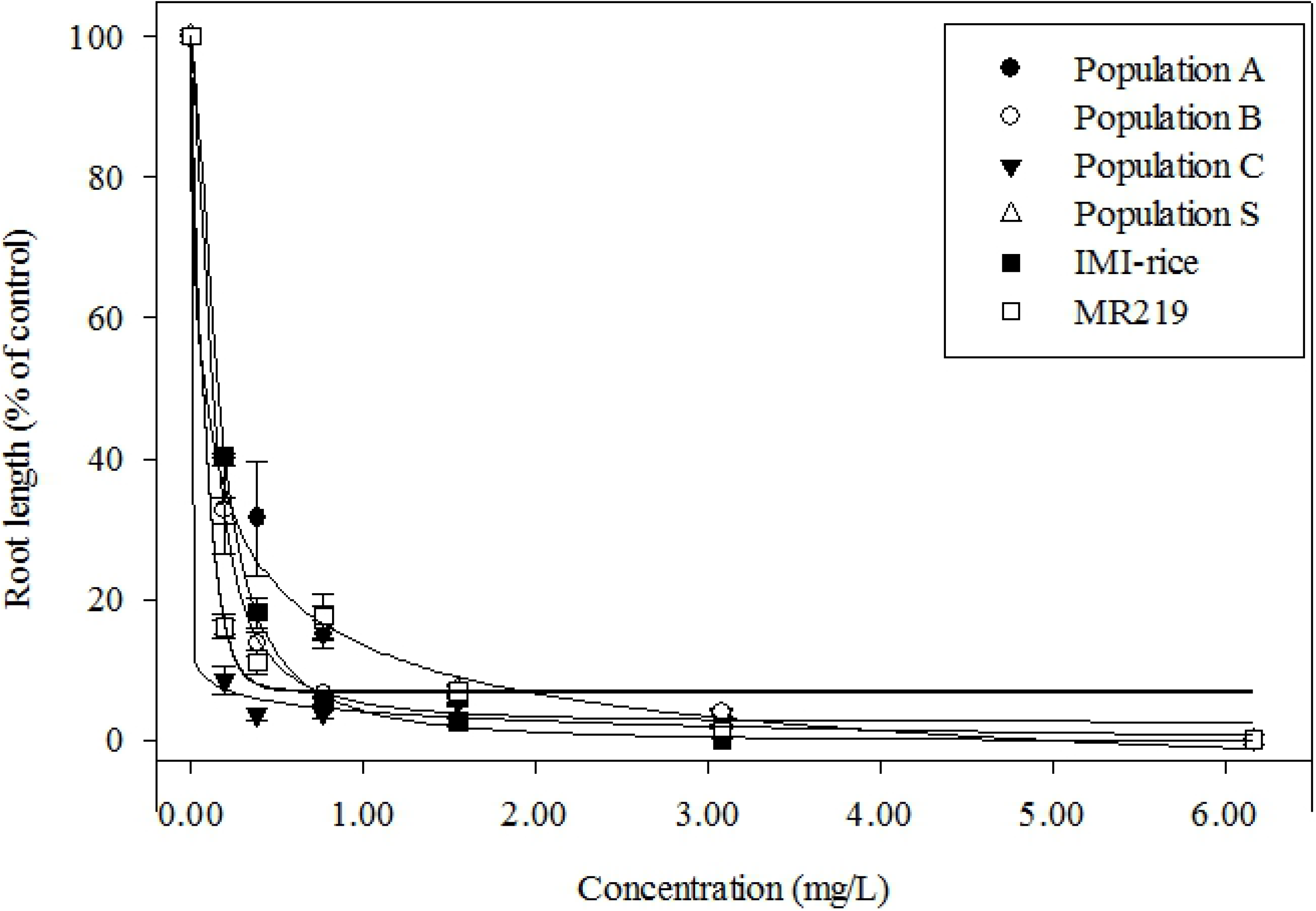
Root length (% of control) of weedy rice populations and rice varieties 14 days after treatment in seed bioassay.

**Fig 4.**
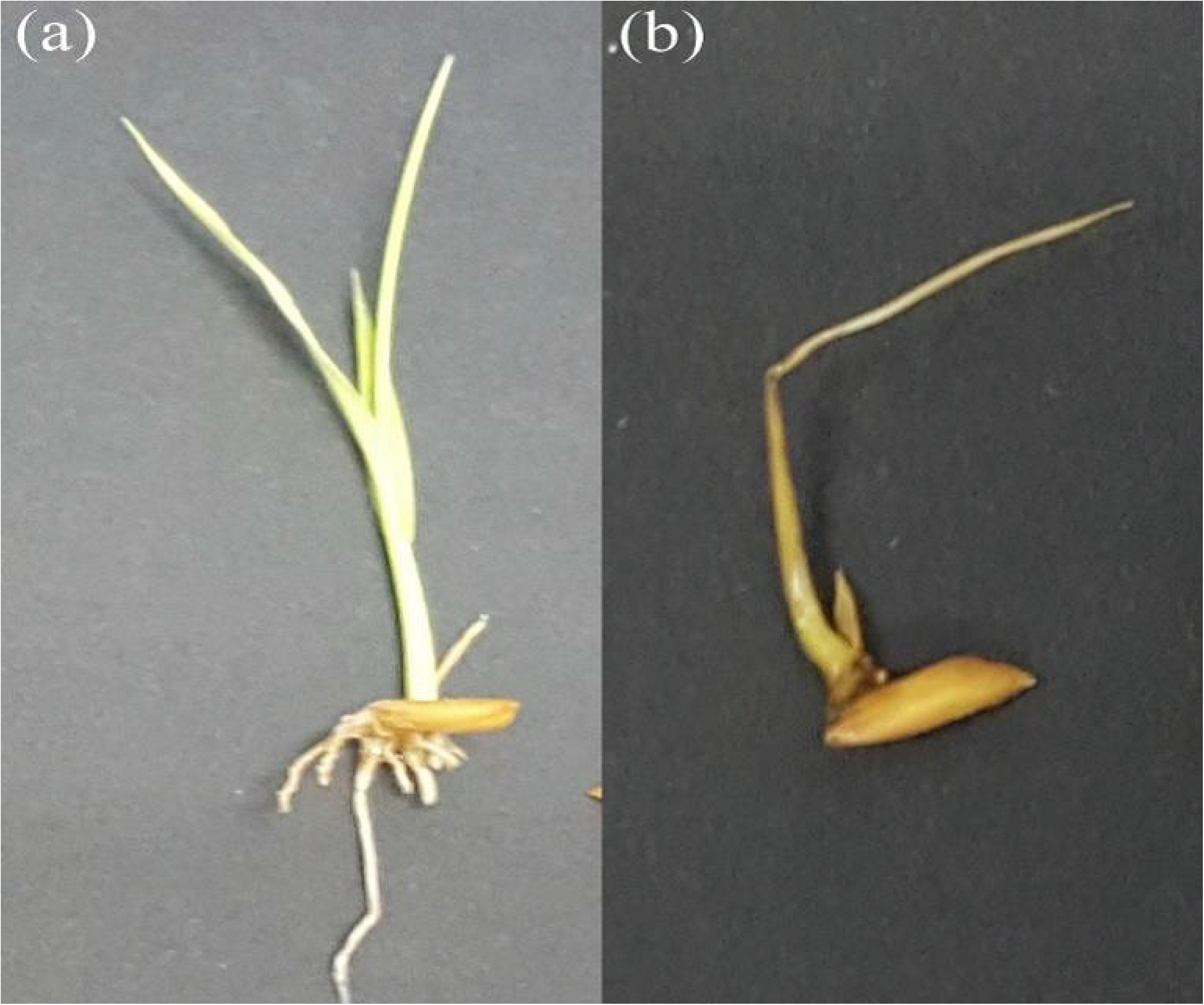
The desiccation sign on seedling shoot of weedy rice populations and rice varieties treated with IMI-herbicides (b) in comparison with the untreated control (a)

**Fig 5.**
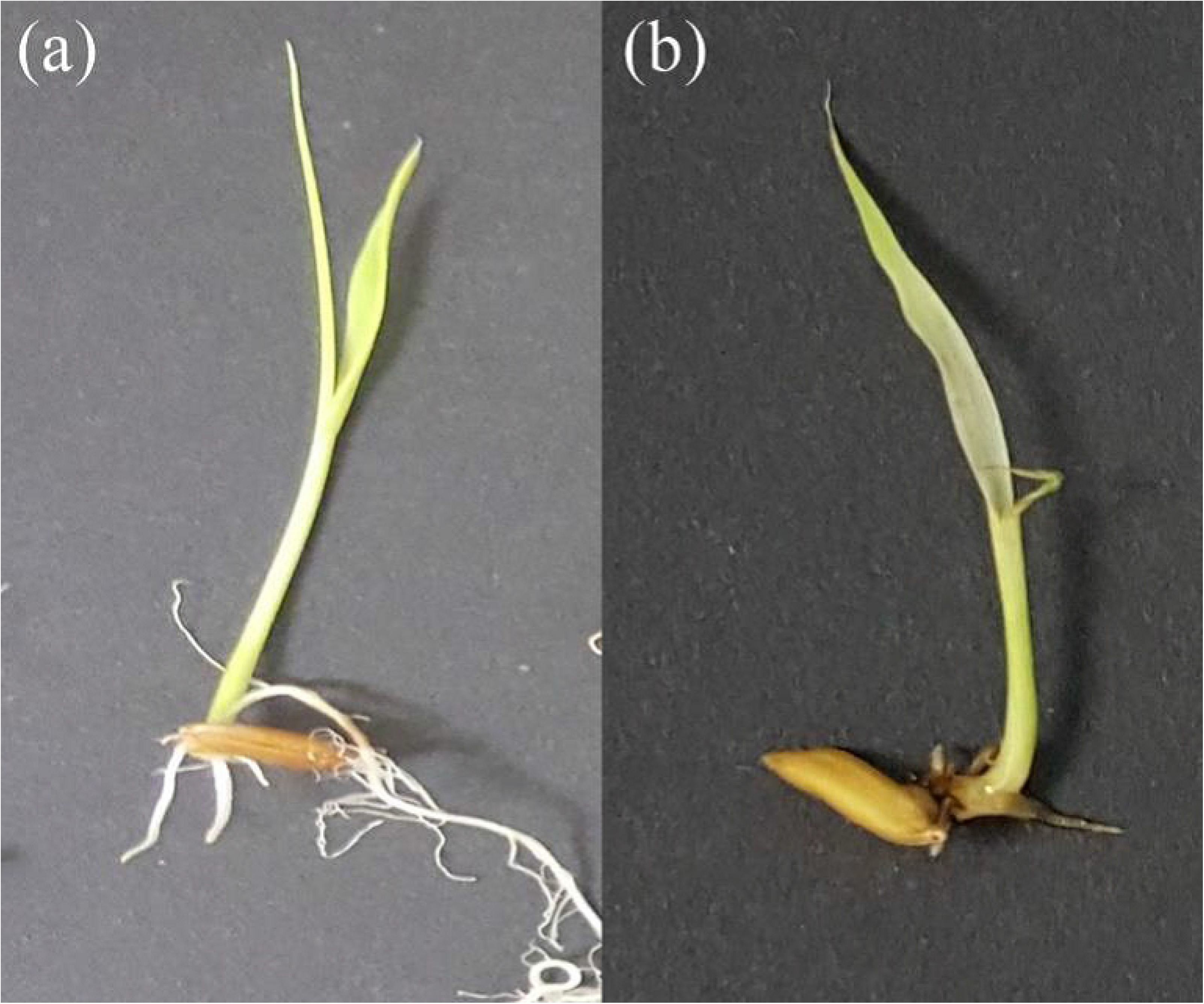
The blackish color on seedling root of weedy rice populations and rice varieties treated with IMI-herbicides (b) in comparison with the untreated control (a)

The effect of IMI-herbicides on shoot length was far more severe than the root length for all weedy rice populations and MR219 rice variety. Nonetheless for IMI-rice, the root was more sensitive to IMI-herbicides than the shoot. Overall, nor shoot neither root growth was recorded at the concentration of 8 times higher than the recommended rate for all populations. Similar with germination rate, both shoot growth and root growth for populations A and B were completely inhibited at the concentration of 8 times higher than the recommended rate (6.16 mg/L). However, the inhibition of shoot and root growth for populations A and B was recorded in different magnitude, where the shoot inhibition ranged from 86-100% while the root inhibition recorded for both populations ranged from 68-100% (Table 1). For population C, the shoots were 100% inhibited at the recommended rate (0.77 mg/L) while the roots managed to elongate until 4-times recommended rate concentration (3.08 mg/L). Similar growth pattern was recorded for S population and rice variety MR219 where the shoots were fully inhibited at only half recommended concentration (0.39 mg/L) while the roots were capable of growing up to 4-times recommended rate concentration (3.08 mg/L). In contrast, the shoots of IMI-rice were still 16% growing at the dose where the roots of the same variety were completely inhibited (3.08 mg/L).

### Whole-plant dose response

Weedy rice populations and rice varieties showed distinguish sensitivity to IMI-herbicides at 21 days after treatment (DAT), with slightly different resistance level against seed bioassay (Table 1), as well as between LD_50_ and GR_50_ (Table 2). The recommended rate of the IMI-herbicides was 154 g a.i/ha. Similar to seed bioassay, IMI-rice variety was the most resistant to IMI-herbicides. It was also noticed that, with the exception of population C, the LD_50_ value resistance level (RI) of all R populations and IMI-rice variety to IMI-herbicides at whole-plant level was lower comparable to LD_50_ RI values in the seed bioassay. Based on the LD_50_ value, IMI-rice displayed moderate level of resistance, while A and B populations exhibited low resistance to IMI-herbicides at whole-plant stage. Both MR219 and S population were equally sensitive. Population C however, exhibited low LD_50_ resistance level at this stage, different than non resistance level expressed in GR_50_ and LD_50_ in seed bioassay (sensitive to IMI-herbicides).

**Table 2.** Analysis of log-logistic of the IMI-herbicides dose needed to exhibit 50% mortality (LD_50_) and to reduce 50% of the shoot dry weight (GR_50_) and the resistance index (RI) of weedy rice populations and rice varieties in whole-plant dose response. Standard errors are in parentheses.

| Populations | LD<sub>50</sub><br>(g a.i/h) | RI<br>(LD<sub>50</sub> R /<br>LD<sub>50</sub> S) | LD<sub>50</sub><br>resistance<br>level | GR<sub>50</sub><br>(g a.i/h) | RI<br>(GR<sub>50</sub> R /<br>GR<sub>50</sub> S) | GR<sub>50</sub><br>resistance<br>level |
| --- | --- | --- | --- | --- | --- | --- |
| A | 216.26<br>(7.22) | 6.43 | Low | 154.91<br>(4.01) | 6.35 | Low |
| B | 86.44<br>(0.72) | 2.57 | Low | 128.97<br>(5.13) | 5.29 | Low |
| C | 121.51<br>(2.94) | 3.61 | Low | 42.61<br>(2.27) | 1.75 | Sensitive |
| S | 33.63<br>(0) | - | Sensitive | 24.40<br>(0) | - | Sensitive |
| IMI-rice | 263.58<br>(3.47) | 7.84 | Moderate | 191.12<br>(7.67) | 7.83 | Moderate |
| MR219 | 34.59<br>(0) | 1.03 | Sensitive | 25.15<br>(0) | 1.03 | Sensitive |

Fig 6 and Fig 7 illustrate a reduction in survival rates and shoot dry weight for all weedy rice populations and rice varieties, respectively, with the exposure of increasing herbicide application rates. No populations survived at the rate of 1200 g a.i/ha (8 times higher than the RR), similar with shoot biomass where 100% of plant shoots were reduced at this concentration. The non IMI-rice MR219 variety can be considered as a positive susceptible control in this experiment. Meanwhile, with the highest LD_50_ and GR_50_ values, the MR220CL2 IMI-rice can be acknowledged as a positive resistant control.

**Fig 6.**
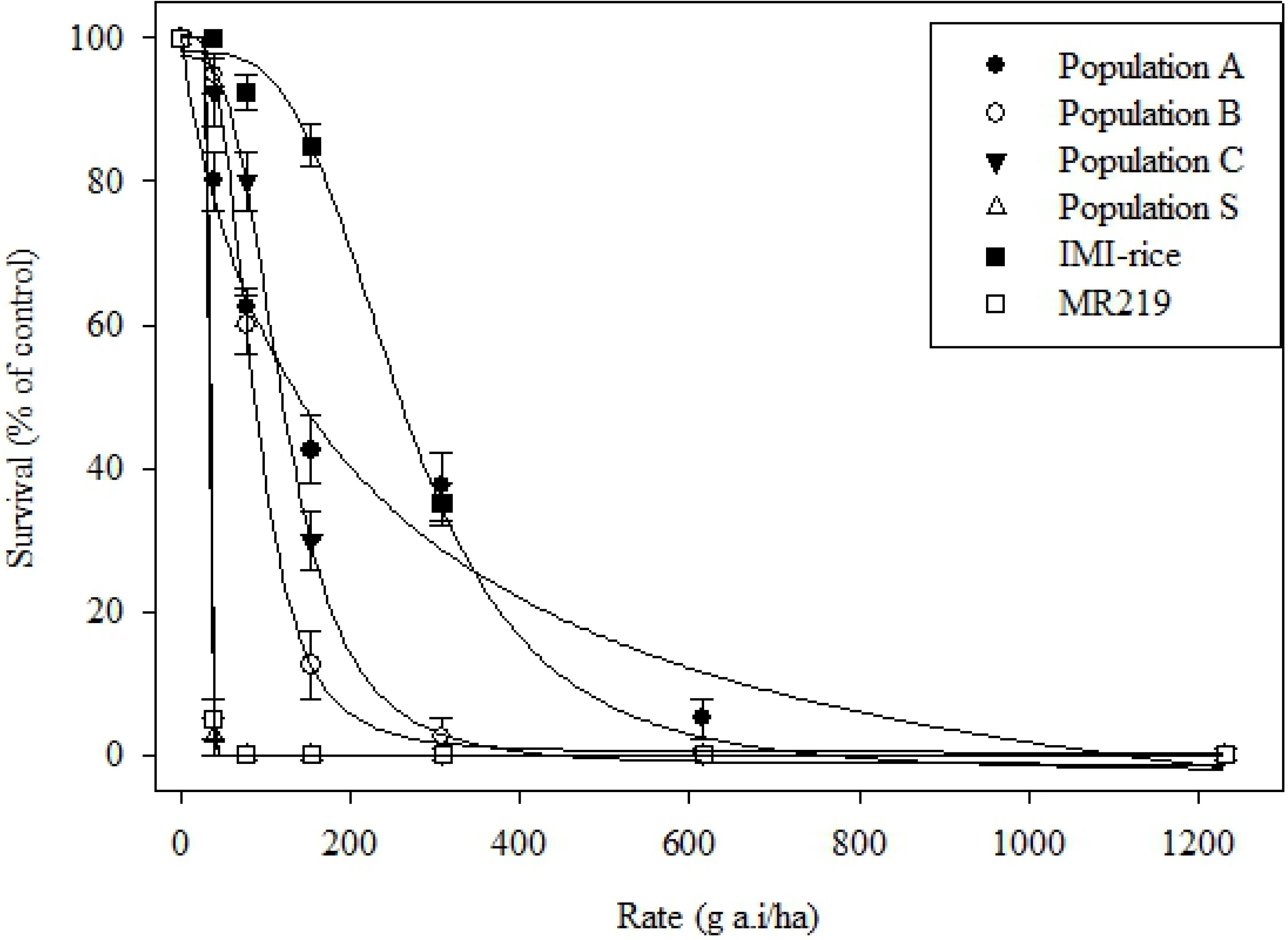
Survival (% of control) of weedy rice populations and rice varieties 21 days after treatment in whole-plant dose response experiment.

**Fig 7.**
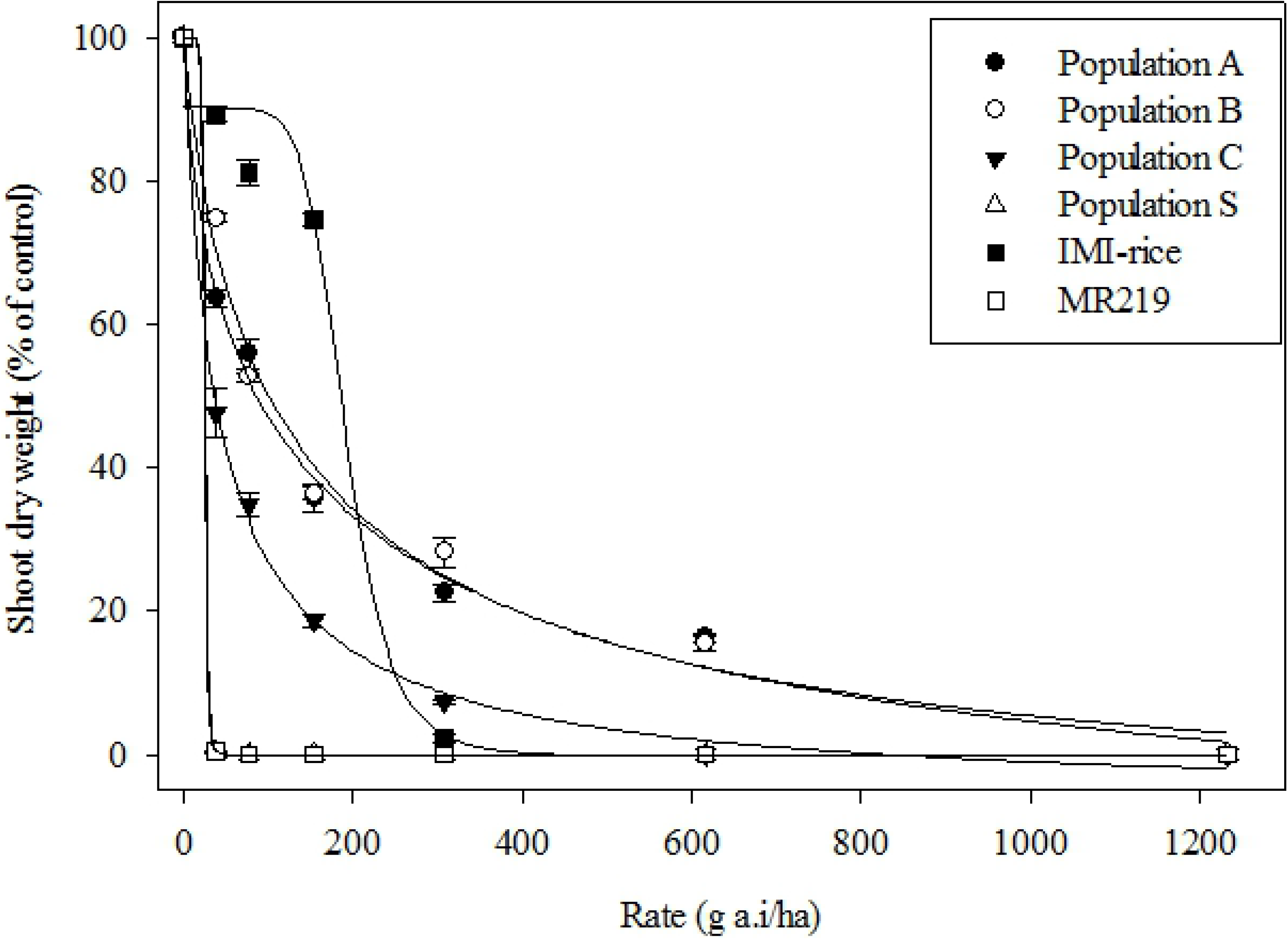
Shoot dry weight (% of control) of weedy rice populations and rice varieties 21 days after treatment in whole-plant dose response experiment.

### Amplification and sequencing of the AHAS gene fragment

The primer pairs yielded the full-length 1884 base pair (bp) AHAS gene of *Oryza* sp. The result from the BLAST showed that the sequenced AHAS gene fragments were identical with the previously characterized *Oryza* AHAS gene available in the GenBank database with 99.8% similarity. The amplified AHAS sequences have been deposited in GenBank (Accession number MN268688 for herbicide-resistant weedy rice and MN268687 for herbicide-susceptible weedy rice). Table 3 shows the nucleotide polymorphisms and amino acid substitutions observed in AHAS gene sequences of R and S weedy rice populations, rice varieties, and *Arabidopsis thaliana*. Four mutations were observed at amino acid positions 93, 557, 653, and 669, where one of these was silent mutation (position 557). The sequenced AHAS gene for R populations of weedy rice was found to have 99% similarity with S weedy rice, while the AHAS gene sequence for R weedy rice was 100% identical to the AHAS coding sequence of IMI-rice variety. A single nucleotide polymorphism (SNP) at amino acid position 653 was revealed in DNA sequence analysis of R weedy rice in relative to S weedy rice, where G was substituted to A at the second base of the same codon in position 653 that resulted in a substitution of amino acid Serine (AGT) by Asparagine (AAT). Interestingly, the similar SNP was observed in IMI-rice AHAS gene when compared to S weedy rice. Meanwhile, the susceptible rice variety MR219 revealed no mutation at Ser_653_ codon, similar to S weedy rice. The presence of AHAS Ser-653-Asn substitution was confirmed in all survived R populations of weedy rice (populations A, B, and C) and IMI-rice with DNA sequencing for at least three individuals, as described by Tardif et al. [22].

**Table 3.**
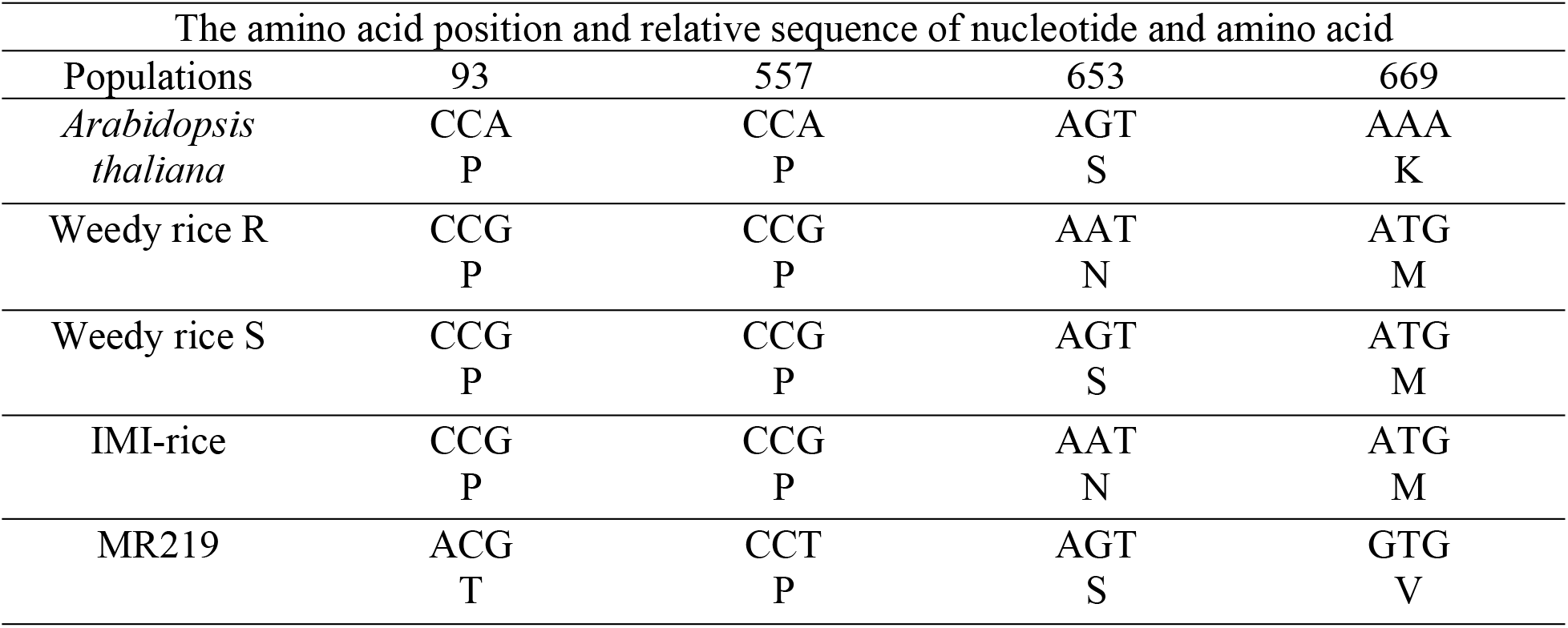
Polymorphisms in nucleotide and amino acid substitutions discovered in the AHAS sequences of *Arabidopsis thaliana*, weedy rice populations, and rice varieties

| The amino acid position and relative sequence of nucleotide and amino acid |  |  |  |  |
| --- | --- | --- | --- | --- |
| Populations | 93 | 557 | 653 | 669 |
| *Arabidopsis thaliana* | CCA<br>P | CCA<br>P | AGT<br>S | AAA<br>K |
| Weedy rice R | CCG<br>P | CCG<br>P | AAT<br>N | ATG<br>M |
| Weedy rice S | CCG<br>P | CCG<br>P | AGT<br>S | ATG<br>M |
| IMI-rice | CCG<br>P | CCG<br>P | AAT<br>N | ATG<br>M |
| MR219 | ACG<br>T | CCT<br>P | AGT<br>S | GTG<br>V |

### *In vitro* AHAS assay

Fig 8 and Fig 9 indicate that the AHAS enzyme activity of all populations following the protein-herbicide incubation decreased with the increase of herbicide concentration. Overall, the AHAS *in vitro* assay for imazapic showed that population A, population B, and IMI-rice were much less sensitive to the herbicide when compared to weedy rice population C, S, and MR219. Meanwhile, referring to the log-logistic curve for imazapyr AHAS inhibition assay, it was clear that all R populations of weedy rice (A, B, and C) and IMI-rice contained AHAS enzyme that is the least sensitive to the IMI-herbicides, while S weedy rice and MR219 were equally susceptible. The I_50_ values for weedy rice A and B populations when treated with different concentrations of imazapic exhibited 605- and 186-fold greater than the S weedy rice, respectively (Table 4). As expected, IMI-rice recorded the highest resistance index (RI) value following imazapic treatment (755.17) compared to the other populations. Meanwhile, the AHAS activity for population C, S, and MR219 rice variety in response to imazapic application showed that the population was highly sensitive to the herbicide. The I_50_ values obtained from this inhibition assay show that population C possessed similar sensitivity level with S weedy rice (0.0012) while, MR219 rice variety was slightly less sensitive to imazapic when compared to S (I_50_ = 0.0017). According to the RI values obtained from this experiment, the populations can be ranked from most resistant to least resistant to imazapic herbicide as follows: IMI-rice > A > B > MR219 > C = S.

**Table 4:** Analysis of log-logistic of the IMI-herbicides dose needed to exhibit 50% inhibition of AHAS activity (I_50_) and the resistance index (RI) of weedy rice populations and rice varieties

| IMI-herbicides | Populations | AHAS activity ($\mu\text{mol}$ acetoin formed/mg protein/h) | $I_{50}$ | RI ($I_{50}$ R/$I_{50}$ S) |
| --- | --- | --- | --- | --- |
| Imazapic | A | 68.83 | 0.7267 | 605.58 |
|  | B | 62.55 | 0.2232 | 186.00 |
|  | C | 65.27 | 0.0012 | 1.0 |
|  | S | 24.22 | 0.0012 | - |
|  | IMI-rice | 71.65 | 0.9062 | 755.17 |
|  | MR219 | 26.17 | 0.0017 | 1.42 |
| Imazapyr | A | 65.45 | 0.3352 | 134.08 |
|  | B | 66.18 | 0.2450 | 77.16 |
|  | C | 63.28 | 0.2398 | 95.92 |
|  | S | 22.74 | 0.0025 | - |
|  | IMI-rice | 78.65 | 0.8697 | 347.88 |
|  | MR219 | 22.98 | 0.0017 | 0.68 |

**Fig 8.**
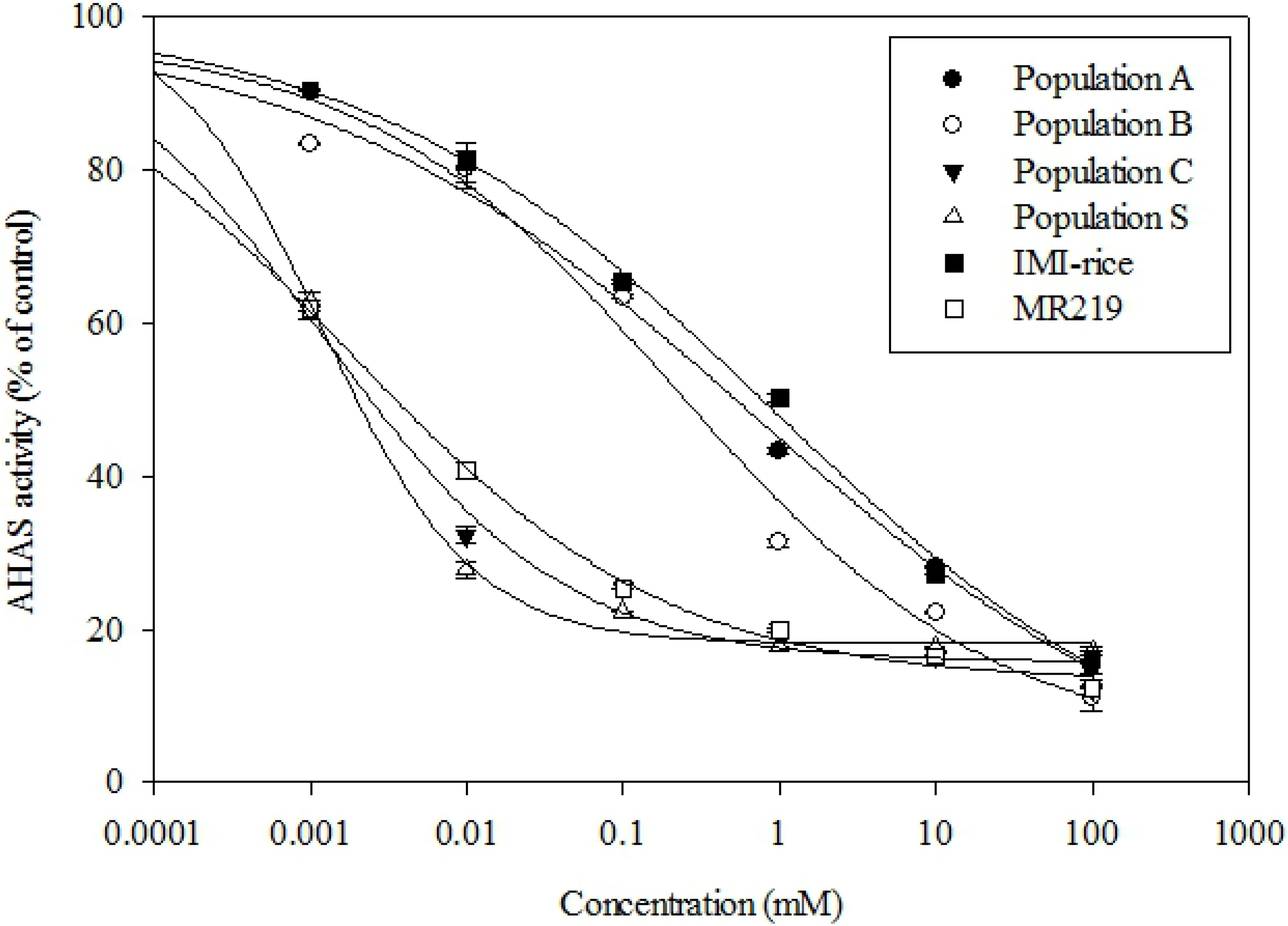
*In vitro* inhibition of AHAS activity by imazapic for weedy rice populations and rice varieties. Data were averaged from three extractions per individual plant of the population and each was assayed four times with standard error of the mean.

**Fig 9.**
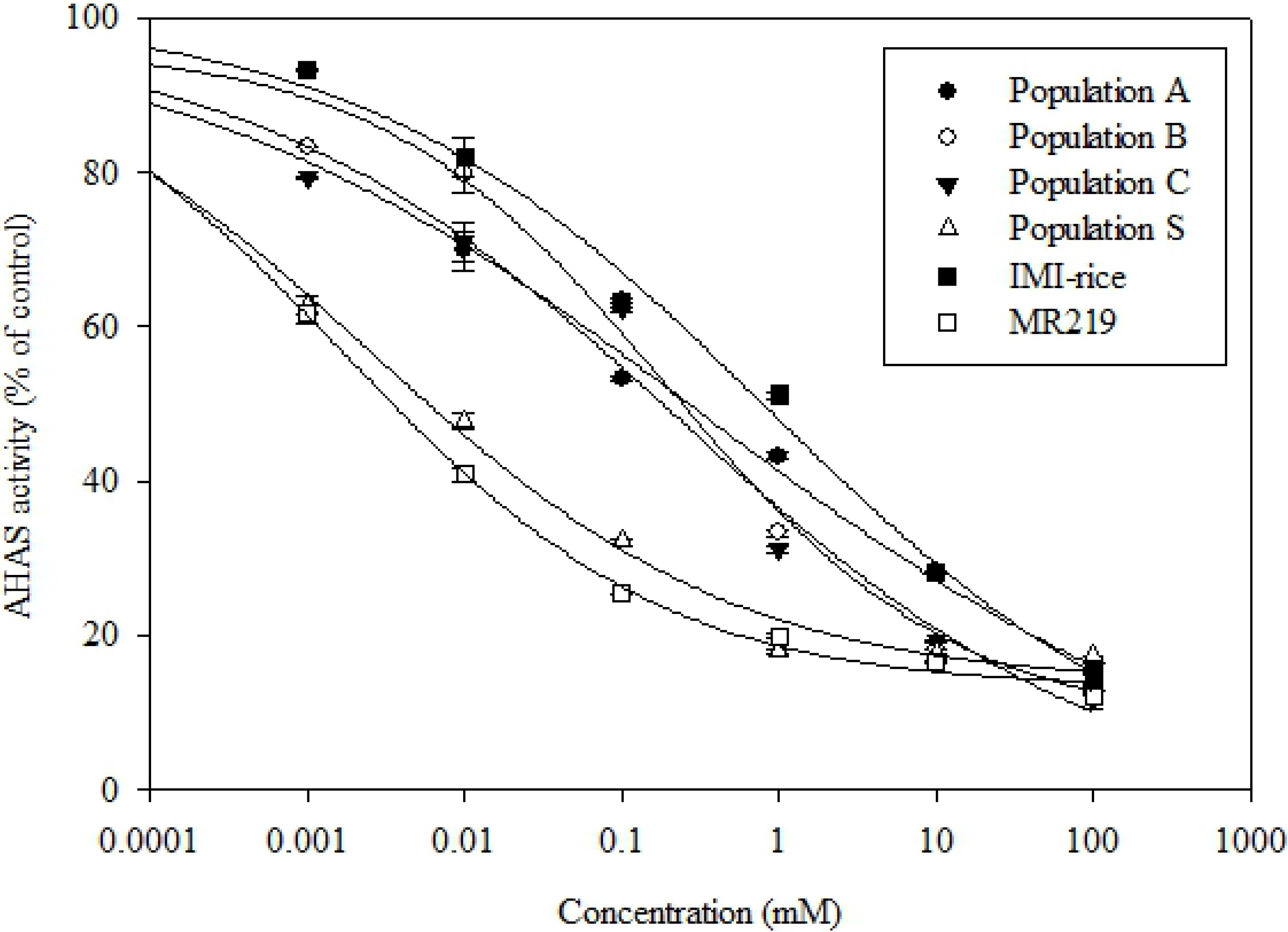
*In vitro* inhibition of AHAS activity by imazapyr for weedy rice populations and rice varieties. Data were averaged from three extractions per individual plant of the population and each was assayed four times with standard error of the mean.

In imazapyr inhibition assay, the similar populations of weedy rice (A, B, and C) were 134, 77, and 95 times more resistant to imazapyr when compared to S weedy rice, respectively. Lower rate of imazapyr was needed to affect half of IMI-rice AHAS activity in comparison with imazapic, although IMI-rice remained exhibiting the highest imazapyr I_50_ RI value over the other populations. Meanwhile, the I_50_ for MR219 was 0.0012, indicated that the population was more sensitive to imazapyr compared to S weedy rice (I_50_ = 0.0025). The ranking of most resistant to least resistant populations of weedy rice and rice varieties to imazapyr herbicide is as follows: IMI-rice > A > C > B > S > MR219. Evidently, population C was also observed less sensitive to imazapyr than imazapic at the enzyme level, which supported slightly higher resistance level in this weedy rice population in whole-plant dose response (Table 2) over the seed bioassay (Table 1).

## Discussion

A screening for herbicide resistance in weeds using petri dish seed bioassay provides a reliable and simple method to evaluate the imidazolinone-resistant gene in weed population [23]. Results from seed bioassay indicate that the continuous exposure of the pre-germinated seeds to the IMI-herbicides has had a significant impact on the seeds growth. The variability of sensitivity to IMI-herbicides among weedy rice populations in petri dish seed bioassay have also been observed in the USA [3,24], Greece [17,25], and Italy [26]. Zhou et al. [27] suggested that AHAS inhibition can be achieved at a very low herbicide concentration, similar with the data obtained for those sensitive populations in this experiment (C, S, and MR219), where up to 99% of inhibition was recorded at only 0.25 times recommended concentration. Because of the ability of the AHAS enzyme to be inhibited at low concentration, Zhou et al. [27] also proposed that this is the reason AHAS becomes the herbicide target.

The seed bioassay results also clearly stipulates that in all weedy rice populations, shoot has been recorded to have higher sensitivity towards IMI-herbicides compared to root. Shivrain et al. [3] also discovered similar finding where, the shoots from US weedy rice accession that had been treated with different concentrations of imazethapyr (also another IMI-herbicide rice package) were more sensitive to the herbicide than the roots. This is not true for the IMI-rice variety nonetheless, where roots growth were slightly more inhibited by the IMI-herbicides, rendering for more investigation on the differential effect of IMI-herbicides on weedy rice and IMI-rice. Roots and shoots are actively dividing meristematic tissues where AHAS is primarily expressed [7]. The biosynthesis of branched-chain amino acids valine, leucine, and isoleucine mainly takes place in young tissues (since these branched-chain amino acids themselves are the crucial components in the development of new cells), making the first injury symptoms appeared after the herbicide inhibition is at the root and shoot growth [27]. As a consequence of IMI-herbicides treatment, seedling shoots also displayed desiccation signs while the inhibited hypocotyl became blackish and eventually led to the plant death, as observed in this study (Figs 4 and 5).

In whole-plant dose response, all weedy rice populations expressed low resistance level towards IMI-herbicides. These results were identical with the whole-plant dose response experiment conducted in another IMI-herbicides adopting country Brazil, where low resistance level was reported in their weedy rice populations treated with similar IMI-herbicides premix of imazapic+imazapyr (weedy rice RI index ranged from 3.5 to 4.5) [28]. Interestingly, population C had a different resistance pattern than the other resistant weedy rice populations and rice varieties at whole-plant level. From the LD_50_ value, population C exhibited low resistance level, contradicted to the sensitive response in seed bioassay. This leaves a question on the misapplication time of IMI-herbicides rice package in the affected area. This was further supported in our recent survey with rice growers across all 10 rice granaries in Peninsular Malaysia, where several farmers practicing IMI-herbicides rice package did admit that they delay the application of IMI-herbicides to 14-day after sowing (Ahmad-Hamdani, pers. comm.), which might have triggered resistance evolution to imazapyr in weedy rice population C. Noticeably, the GR_50_ RI index in population C also was already 1.75 (nearly 2 to become resistance at low level). This was further proved by the I_50_ value, in which population C had more sensitive enzyme response towards imazapic than imazapyr (Table 4). Thus in a matter of time, if inappropriate application time of IMI-herbicides is continuously practiced, it is likely for the imazapic-imazapyr resistance pattern to shift and escalate in some weedy rice populations.

The IMI-rice recorded the highest LD_50_ value, possessing moderate resistance level to IMI-herbicides (Table 2). Interestingly, even though this herbicide tolerant rice variety exhibited higher tolerance/resistance level to the IMI-herbicides over the weedy rice populations, a significant reduction in survival rate and shoot dry weight was recorded at a concentration of 2 times higher than the recommended rate (308 g a.i/ha) (Figs 6 and 7). This finding shows that although IMI-rice moderately survived the IMI-herbicides, the growth was still impeded to a certain degree. Thus, it is important for farmers to follow the spray recommendation rate and IMI-rice growth condition in order to reduce the risk of growth retardation or fitness penalty/cost in IMI-rice to ensure high rice yield.

The IMI-herbicides OnDuty™ WG premix comprise of imazapic (52.5%) and imazapyr (17.5%) as active ingredients. Sartori et al. [29] opined that the application of imazapic as pre-emergence was effective in controlling weedy rice. Similar outcome was reported by Washburn and Barnes [30] where pre-emergence application of imazapic was effective to control grass weeds compared to post-emergence application. Mangold et al. [31] also reported that imazapic provides better control to downy brome seedlings when applied immediately after emergence than applying it to older seedlings. Meanwhile, the effectiveness of imazapyr is increased when applied as a post-emergence herbicide even though it can be applied as both pre- and post-emergence [32].

As stated previously, with the content percentage of 52.5% for imazapic and 17.5% for imazapyr, the OnDuty™ WG premix consists higher amount of pre-emergence herbicide imazapic than post-emergence imazapyr. This is the reason for the application recommendation of OnDuty™ WG herbicide in commercial IMI-rice fields, where it must be applied as pre-emergence or early post-emergence to the plants (preferably 0-5-day after sowing, and not exceed 7-day after sowing or 1-2-leaf stage). Seed bioassay was conducted to evaluate the pre-emergence activity of the IMI-herbicides premix to weedy rice populations and rice varieties, while whole-plant dose response experiment allowed us to confirm resistance of the populations with post-emergence activity [33]. Hence, both experiments aimed to evaluate the occurrence and level of resistance of weedy rice populations and rice varieties to imazapic and imazapyr that best applied as pre- and early post-emergence herbicides, respectively. In seed bioassay, weedy rice populations possessed different level of resistance to IMI-herbicides (high, moderate, sensitive) (Table 1). Conversely, all of these populations of weedy rice were found to have low resistance level to IMI-herbicides in whole-plant dose response experiment (Table 2). This results clearly define the cross-resistance pattern of weedy rice populations and rice varieties towards imazapic and imazapyr.

The PCR amplification of all plant samples from weedy rice populations and rice varieties managed to successfully amplify the entire AHAS gene fragment (1884 bp) that contain all six conserved regions as described by Merotto et al. [21]. The conserved regions C, A, D, F, B, and E were also known as domains, where mutations endowing herbicide-resistance have been previously found in these particular regions. The comparison of AHAS sequences for all plant samples of weedy rice populations and rice varieties indicated 99% similarity. This proves that the AHAS gene sequences are highly conserved in *Oryza* spp. either in weedy rice or cultivated rice. The sequenced AHAS gene fragment of IMI-susceptible rice variety MR219 and S weedy rice was 99% identical at nucleotide level in the AHAS coding region (Table 3). Mutations at positions 93, 557, and 669 lied outside of the conserved domains.

The amino acid substitution in position 93 and position 669, resulted in Pro_93_Thr and Val_669_Met, respectively was evident in all weedy rice populations and IMI-rice. Shivrain et al. [3] recorded similar Pro-93-Thr and Val-669-Met mutations in weedy rice accessions collected in Arkansas, USA relative to IMI-herbicide susceptible local rice cultivar Bengal. Both Pro_93_ and Val_669_ that existed in Bengal rice cultivar were substituted to Thr_93_ and Met_669_ in weedy rice populations and IMI-rice cultivar (Table 3). In addition, Sales et al. [34] also recorded similar Val-669-Met substitution in the USA resistant red rice accessions Lon-2 and Ran-4 when compared to Bengal rice cultivar. Both findings were consistent with the result obtained in this study where the amino acid Val_669_ in weedy rice populations was altered to Met_669_ in R and S weedy rice, and also IMI-rice.

Sales et al. [34] suggested that the Val_669_Met substitution in AHAS gene of resistant USA red rice accessions might contribute to the insensitivity of the enzyme towards IMI-herbicides. The crystal 3D structure of *Arabidopsis thaliana* AHAS (*At*AHAS) showed that the enzyme has four identical subunits [35]. There are three domains associated in each of these subunits namely α (residues 86-280), β (residues 281-451), γ (residues 463-639), and a C-terminal tail (residues 646-668) [7,35]. Val_669_ is located near to the C-terminal tail of the AHAS enzyme structure. Until the study was reported by Sales et al. [34], the crystal structure of the C-terminal tail in plant AHAS has yet to be discovered. Thus, initial presumption of the Val_669_ relation to the enzyme insensitivity to the herbicide was made by referring to the crystal structure of C-terminal tail in fungal, *Saccharomyces cerevisiae* AHAS (*Sc*AHAS). However, recently, Garcia et al. [36] revealed that the plant AHAS crystal structure is significantly different with the fungal AHAS. The absence and presence of herbicides give an impressive effect to the fungal enzyme. The effects and changes include the decreased in volume of accessible space around the active site, position of cofactor FAD and ThDP in active site, and ordering of the “capping region” or “mobile loop” and C-terminal arm, which are influential in herbicide-binding. Nonetheless, Garcia et al. [36] found that the overall amino acid sequence of *Sc*AHAS and *At*AHAS is substantially different to each other, especially in the capping region (the “mobile loop” and the C-terminal arm of residues L568-R583 and H646-R667) which then proved that in plant AHAS, the herbicide-binding site is highly flexible and adaptable depending on the specific identity of the herbicide that binds, which made them contrast with fungal AHAS.

Thus, considering that this mutation positions outside the binding domains of AHAS inhibitors, it is not convincingly relevant to relate the Val_669_Met substitution to the reduced sensitivity of the enzyme to AHAS herbicides until further study on the enzyme structure associated to this amino acid substitution is conducted. The results obtained in this experiment also corroborate that the Val_669_Met substitution is not a resistant-bearing mutation since the substitution occurred in both populations of R and S weedy rice (Table 3). In seed bioassay and whole-plant dose response, the survived R populations of weedy rice was proved to have low to high resistance level to the IMI-herbicides, while the Val_669_Met mutation-bearing S weedy rice was extremely susceptible to the herbicide.

Rajguru et al. [12] stated that the weedy species of rice and cultivated rice are expected to have further genetic differences in other loci that clarify the phenotypic plasticity of the weedy species, especially in the unamplified regions. Although *Oryza* spp. generally exhibit low genetic variation within a population because of its self-pollinated feature, genetic variation can occur widely among the populations [3,37]. For instance, the occurrence of pollen-mediated gene flow and hybridization between the cultivated rice and weedy rice has been evidenced in several studies [12,34,38,39,40]. The gene flow and hybridization events that occur among *Oryza* spp. also have contributed to the genetic variation of weedy species of rice in Malaysia [41]. Song et al. [41] recommended that the modern elite rice cultivar is influential in the genetic evolution of Malaysian weedy rice. The recent morphological study of Malaysian weedy rice also supports the perception that the origin of Malaysian weedy rice was mainly contributed by the wild species of *Oryza* and the modern-bred elite cultivars [42]. The 99% identity of AHAS gene between the weedy rice S and MR219 in this study represented a close resemblance of Malaysia weedy rice to its local cultivated variety of rice.

It is widely known that the G to A substitution in amino acid position 653 from Serine to Asparagine is responsible for herbicide resistance/tolerance in rice CPS technology [43,44]. Malaysia IMI-rice cultivars (MR220CL1 and MR220CL2) were derived from crosses between Malaysia local rice variety MR220 and Louisiana State University (LSU) North America IMI-rice cultivar CL1770 that harbors a Ser-653-Asn mutation [43]. Thus, the presence of Ser-653-Asn mutation in AHAS gene of Malaysian IMI-rice variety was confirmed through this study. The SNP that causes substitution in amino acid 653 in IMI-rice was expected as this is the mechanism of resistance to IMI-herbicides in IMI-resistant rice cultivar, as well as other species of weeds, including weedy rice [12,45]. It has been reported by many researchers that the most common mechanism endowing resistance to AHAS inhibitors in weed species is due to amino acid substitutions within the AHAS gene. To date, a total of 27 amino acid substitutions in AHAS gene at eight conserved positions have been identified to confer resistance to AHAS inhibitors in various weed species [46]. The eight conserved positions conferring resistance to IMI-herbicides are Ala_122_, Pro_197_, Ala_205_, Asp_376_, Arg_377_, Trp_574_, Ser_653_, and Gly_654_ [46, 47]. In this study, a resistance-endowing polymorphism was only observed at one position out of eight conserved positions reported, which is at amino acid position 653.

The Ser-653-Asn mutation is particularly interesting to be discussed because Ser_653_ is located in a highly conserved amino acid domain that has caused field-evolved resistance to AHAS inhibitors among plants [21] and commonly reported in other studies of AHAS-inhibitor resistant species. Tranel and Wright [48] summarized that several single changes in amino acid of AHAS enzyme are capable of converting the herbicide-susceptible plant to a herbicide-resistant plant, and Ser_653_ that located at the carboxyl-terminal of the gene is one of the important resistance-bearing mutation. A Ser-653-Thr substitution was reported in *Amaranthus powellii* S. Wats. that survived the application of imazethapyr and exhibit cross-resistance to atrazine [49]. Additionally, a point mutation resulting in a substitution of Ser_653_ by Asn_653_ was found in resistant plant of *Amaranthus rudis* Sauer [50]. A similar mutation was also reported in imazethapyr-resistance Strawhull hybrid red rice in southern United States, where the same Ser-653-Asn mutation was also observed in the cultivated Clearfield^®^ rice variety [12]. In IMI-resistant red rice found in Southern Brazil, Ser-653-Asp and Ser-653-Asn mutations were found in red rice samples collected from both years of 2006/2007 and 2007/2008 [51]. Similarly, a study conducted by Busconi et al. [52] also found that the Ser-653-Asn mutation was observed in the resistant red rice accessions escaped from Italy Clearfield^®^ rice fields. Subsequently, the same study was conducted to imazamox-resistant red rice populations in Italy where all of the resistant populations possessed a Ser to Asn substitution at locus 653 of the gene [53]. Other than that, a homozygous mutation of Ser-653-Asn was recorded in both northern Greece putative resistant red rice and Clearfield^®^ rice cultivar where, further PCR for ‘Clearfield_®_ allele’ detection proved that the genetic background of putative resistant red rice matched with the Clearfield^®^ rice cultivar [17].

The *in vitro* AHAS assay results in weedy rice populations and rice varieties were equivalent to the results obtained in seed bioassay and whole-plant dose response experiment. Meanwhile, the pattern of resistance conferred by the Ser-653-Asn mutation in Malaysia IMI-rice was slightly different with the other IMI-resistant rice with the same mutation, for example, Avila et al. [54] found that the CL-161 rice variety that harbors Ser-653-Asn mutation was 420 times more resistant to imazethapyr than its relative S weedy rice ecotypes and conventional rice variety Cypress. However, the RI recorded for Malaysia IMI-rice in *in vitro* enzyme assay revealed that the rice variety was 755-fold more resistant to imazapic and 347-fold more resistant to imazapyr, in comparison with S population. The variability of resistance pattern at enzyme level in the populations endowing similar point mutation to IMI-herbicides can be explained by the plant AHAS crystal structure modelling. The difference in chemical structure of AHAS inhibitors, even though from the same family would influence the herbicide binding orientation of the herbicide to the target domain of the enzyme, in which different IMI-herbicide possesses different binding orientation [35,47,55]. The similar reason can possibly explain the variability of the resistance level of weedy rice populations to imazapic and imazapyr in this experiment. The RI index recorded for all the populations varied considerably even though all of the survived plants involved in this present study harboring the same Ser-653-Asn mutation.

To conclude, this study proves cross-resistance pattern to IMI-herbicides imazapic and imazapyr in different weedy rice populations, contributed by insensitive AHAS enzyme, particularly target-site Ser-653-Asn mutation in AHAS gene. Populations A and B expressed higher resistance level to imazapic, while population C exhibited resistance development to imazapyr. This is the first report in AHAS-herbicides resistance mechanism in weedy rice in Malaysia.

